# Nanobody engineering for SARS-CoV-2 neutralization and detection

**DOI:** 10.1101/2022.09.14.507920

**Authors:** Liina Hannula, Suvi Kuivanen, Jonathan Lasham, Ravi Kant, Lauri Kareinen, Mariia Bogacheva, Tomas Strandin, Tarja Sironen, Vivek Sharma, Petri Saviranta, Anja Kipar, Olli Vapalahti, Juha T. Huiskonen, Ilona Rissanen

## Abstract

In response to the ongoing SARS-CoV-2 pandemic, the quest for coronavirus inhibitors has inspired research on a variety of small proteins beyond conventional antibodies, including robust single-domain antibody fragments, ‘nanobodies’. Here, we explore the potential of nanobody engineering in the development of antivirals and diagnostic tools. Through fusion of nanobody domains that target distinct binding sites, we engineered multimodular nanobody constructs that neutralize wild-type SARS-CoV-2 and the Alpha and Delta variants with high potency, with IC50 values up to 50 pM. However, we observed a limitation in the efficacy of multimodular nanobodies against the Beta (B.1.351) and Omicron variants (B.1.1.529), underlining the importance of accounting for viral evolution in the design of biologics. To further explore the applications of nanobody engineering in outbreak management, we present a novel detection assay, based on fusions of nanobodies with fragments of NanoLuc luciferase that can detect sub-nanomolar quantities of the SARS-CoV-2 spike protein in a single step. Our work showcases the potential of nanobody engineering to combat emerging infectious disease.

## Introduction

Antibody-based products comprise some of the most successful diagnostic and therapeutic tools developed for managing the COVID-19 pandemic, ranging from rapid antigen tests for SARS-CoV-2 infection (1, 2) to neutralizing monoclonal antibodies (mAbs) used to treat COVID-19 in individuals at risk of severe disease (2, 3). Neutralizing antibodies against SARS-CoV-2 primarily target the Spike (S) protein (4, 5), a glycoprotein mediating host-cell recognition and viral entry (6). SARS-CoV-2 spikes are homotrimers, with each chain consisting of receptor-binding (S1) and fusogenic (S2) subunits (7). S1 contains the receptor- binding domain (RBD) which mediates binding to the primary cellular receptor of SARS-CoV- 2, angiotensin-converting enzyme 2 (ACE2) (6, 8, 9). Following receptor binding, S2 subunit, a class I fusion protein, is activated by proteolytic cleavage and mediates fusion of the viral and cell membranes, delivering viral RNA to the cytoplasm (8, 10).

Due to its key role in initiating infection, the SARS-CoV-2 spike is the primary target of vaccines (11) and monoclonal antibody therapy (2). The continued efficacy of these powerful approaches is challenged by the emergence of SARS-CoV-2 variants of concern (VOCs) that display multiple amino acid substitutions in the S-protein (12-14). Following the spread of variants Alpha (B.1.1.7), Beta (B.1.351) and Delta (B.1.617.2) (15-20), Omicron (B.1.1.529) has become the dominant circulating variant in 2022, with new Omicron sub-variants still emerging (21). VOC amino acid changes, including E484K found in Beta and Omicron, can reduce neutralization by antibodies raised against the ‘wild-type’ SARS-CoV-2 (titled B.1 or Wuhan-Hu-1) (12, 15, 16, 19, 22). Current efforts to mitigate effects of immune escape on antibody-based COVID-19 countermeasures include use of antibody cocktails (23, 24) and development of new antibody-based products, including camelid single-domain antibody fragments (‘nanobodies’) (25).

In contrast to traditional mAbs, nanobodies are small (∼15 kDa) and offer advantages including nebulized delivery and scalable, cost-effective production in bacterial expression systems (26, 27). During the COVID-19 pandemic, antiviral nanobodies have garnered significant interest, resulting in discovery and structural characterization of several SARS-CoV-2-neutralizing nanobodies (27-37). Furthermore, as nanobodies comprise self-contained modules, they can be engineered into fusion proteins with enhanced properties. Pioneering studies have started to chart the potential of engineered nanobodies as virus inhibitors (38, 39), but diagnostic applications are largely unexplored. While RT-qPCR remains the gold standard for clinical diagnosis, rapid diagnostic tests designed to detect viral antigens with conventional antibodies (40) are extensively applied in nonhospital settings. To our knowledge, nanobodies have not previously been used for viral antigen detection in commercialized assays.

Here, we explore engineering nanobody fusions for enhanced neutralization and development of a novel rapid-antigen assay. Tri-modular fusions of selected nanobodies showed up to a thousandfold enhancement of the *in vitro* neutralization efficiency against wild-type SARS-CoV-2 as compared to the reported efficiencies of constituent nanobodies (28-30, 41). Nanobody fusions were further engineered to produce proof-of-concept for a novel diagnostic assay, which applies nanobodies fused to fragments of a split signal molecule, NanoLuc luciferase (42-45), and allows the detection of picomolar concentrations of SARS-CoV-2 spike protein in a single step. Overall, our study shows the potential for engineered nanobodies as antiviral and diagnostic agents, which we envision can offer affordable and scalable countermeasures during future outbreaks of emerging viral diseases.

## Results

### Structure-guided design of multimodular nanobodies

Inspired by the trimeric structure of the coronaviral spike (Figure 1), we sought to develop an approach for targeting all RBDs simultaneously to enhance SARS-CoV-2 inhibition. Cryogenic electron microscopy (cryo-EM) studies have identified two SARS-CoV-2 RBD conformations, ‘up’ and ‘down’ (46, 47), where putative epitopes on neighboring RBDs are in proximity, within 40 to 74 Å from each other (SI Figure S1). To develop a tripartite binder that could be sterically accommodated within these tight constraints, we selected nanobodies, the smallest antibody-based protein inhibitors, as the base unit for multimodularization.

**Figure 1.**
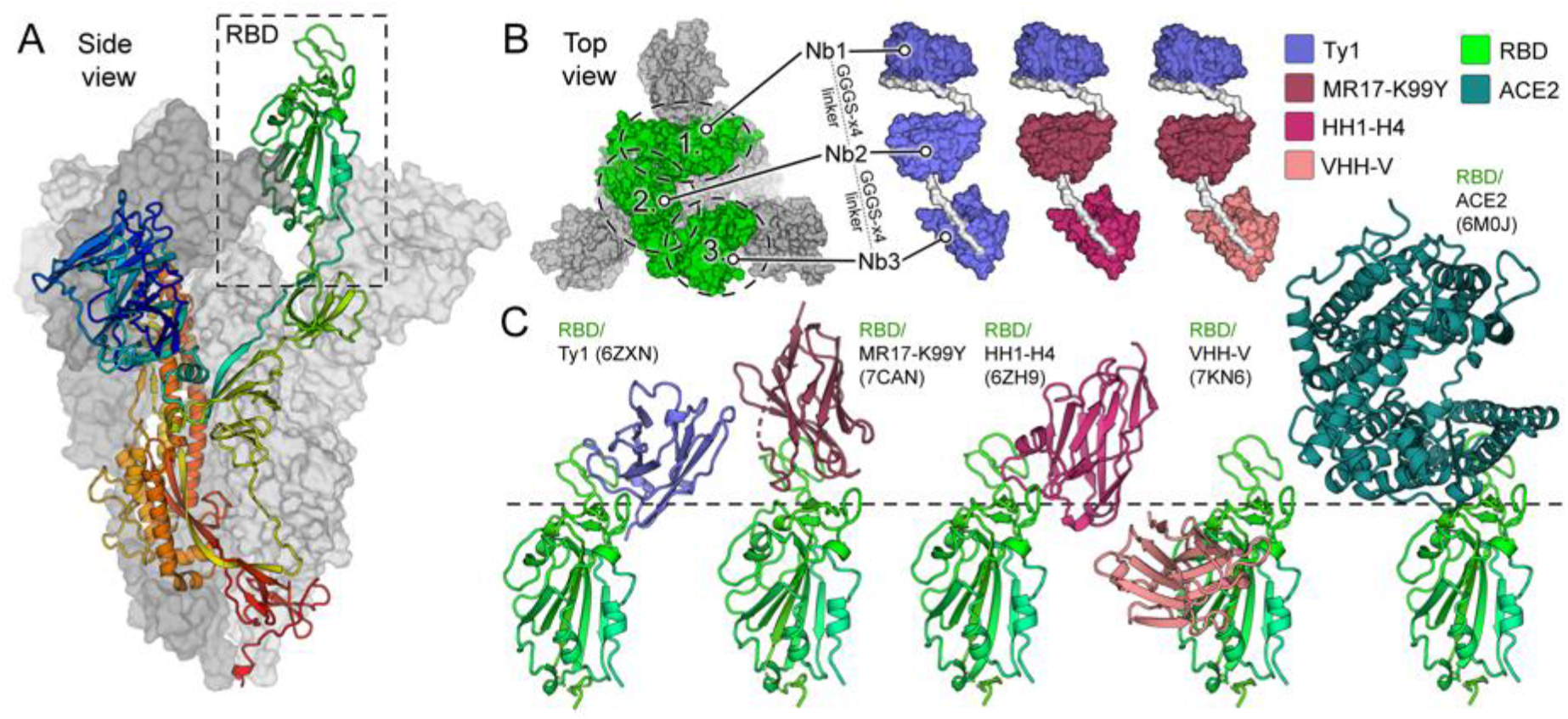
Structure-based design of multimodular nanobodies targeting SARS-CoV-2 S. **A**) Side view of the SARS-CoV-2 S-trimer (28). One trimer subunit, with a receptor-binding domain (RBD), in the ‘up’ conformation, is shown as a cartoon and colored as a rainbow ramped from blue (N-terminus) to red (C-terminus). The two other subunits are shown as surface representation in shades of grey. **B**) Three distinct multimodular nanobodies were designed: (i) tri-Ty1 with three repeats of Ty1 (28) module; (ii) tri-TMH with Ty1, MR-17- K99Y (30) and HH1-H4 (29) modules; and (iii) Tri-TMV with Ty1, MK and VHH-V (41) modules. The nanobody modules are connected by flexible GGGGSx4 linkers. **C**) Each of the nanobody domains (Ty1, MR17-K99Y, HH1-H4, and VHH-V) bind the RBD at a unique angle. Comparison to the structure of RBD bound to the primary host-cell entry receptor of SARS-CoV-2, ACE2 (6, 80) demonstrates that three of the modules bind epitopes proximal to the ACE2-binding site, while VHH-V targets an alternative neutralization epitope.

We designed three trimodular nanobodies (Figure 1B) using the sequences of four previously published monomeric nanobodies, Ty1 (28), H11-H4 (29), MR17-K99Y (30) and VHH V (41), reported to neutralize wild-type SARS-CoV-2 with IC50 (half-maximal inhibitory concentration) values ranging from 38 to 142 nM (SI Table S1). These modules were selected based on two criteria: (i) distinct epitope and angle of binding to RBD (Figure 1C and SI Table S1), (ii) spatial proximity of the epitopes in the context of the SARS-CoV-2 spike, facilitating simultaneous binding of all modules (Figure 1).

Multimodular nanobodies were constructed by fusing three nanobodies together with flexible linkers of twenty amino acids (GGGGSx4) with the aim to improve binding avidity (26, 39, 48-50). Here, construct compositions were selected to test i) the effect of triplicating a single module on SARS-CoV-2 neutralization, and ii) whether the inclusion of variable modules helps mitigate neutralization escape (41). To this end, multimodular constructs tri-Ty1 (comprised of three Ty1 modules (28)), tri-TMH, and tri-TMV (Figure 1B) were generated, with tri-TMH and tri-TMV comprised of **T**y1 (28) and **M**R17-K99Y (30) modules, followed by either a **H**11- H4 (29) or VHH **V** (41) module, respectively.

### Multimodular nanobodies bind variant forms of the RBD

Amino acid changes observed in SARS-CoV-2 VOCs have been linked to escape from antibody-mediated neutralization (15, 16, 51, 52) due to reduced affinity to epitopes with altered amino acids. To determine how RBD amino acid changes K417N, E484K, and N501Y impact multimodular nanobodies, we tested their binding to RBD and spike mutants in an antigen microarray. Nanobodies tri-Ty1, tri-TMH, and tri-TMV were tested against (i) wild-type RBD, (ii-iv) three RBD variants that displayed either K417N, E484K, or N501Y amino acid change, (v) wild-type S1 subunit, and (vi) a S1 subunit displaying amino acid changes K417N, E484K, N501Y, and D614G. Tri- TMH and tri-TMV each contain three distinct nanobody modules targeting a broad range of residues, and therefore were expected to be less sensitive to amino acid changes than tri-Ty1. The results support this hypothesis, showing that binding of tri-Ty1 to RBD was strongly decreased by the E484K change, while this effect was mitigated in tri-TMH and tri-TMV (Figure 2). Amino acid changes K417N or N501Y did not notably alter binding.

**Figure 2.**
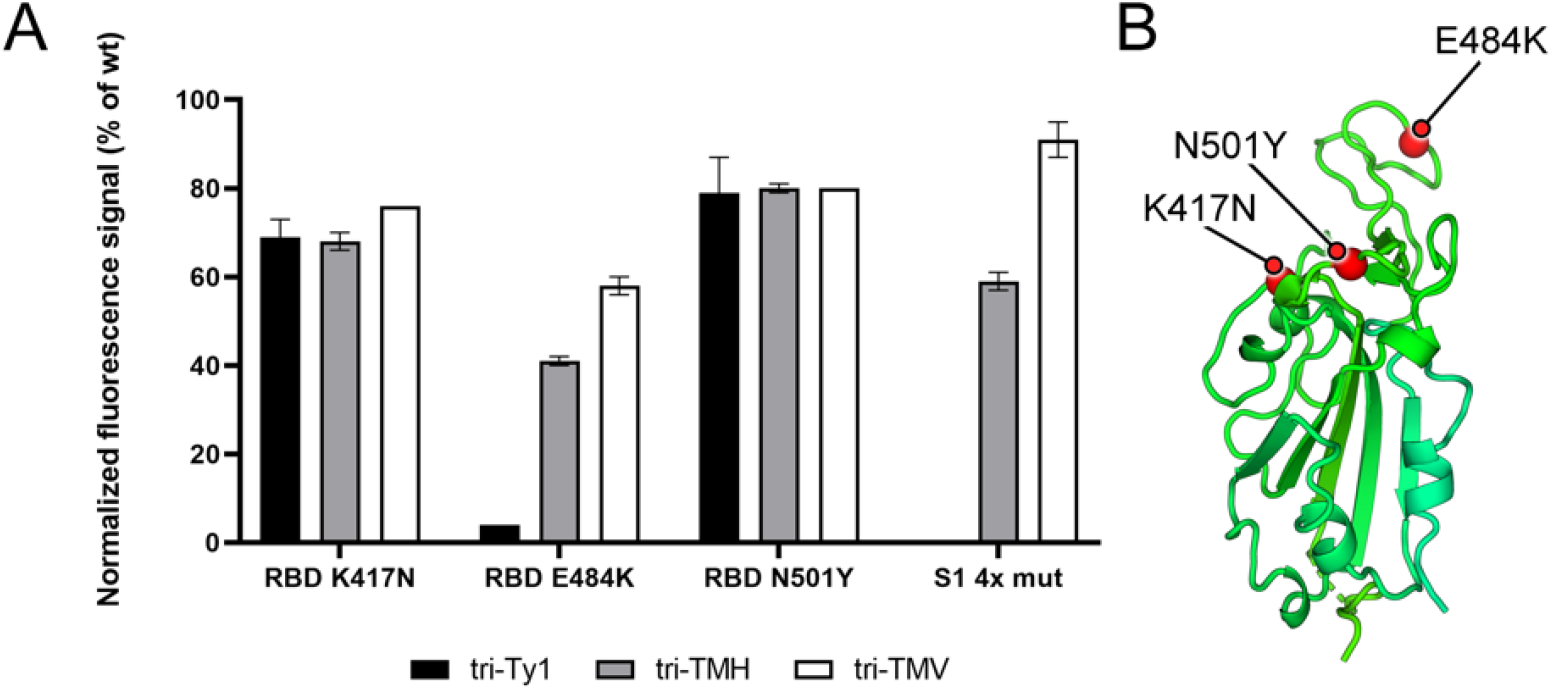
Relative binding strength of trimeric nanobodies to different RBD and Spike S1 domain variants on the antigen microarray. **A**) Three multimodular nanobody constructs were tested in a binding array and were shown to bind different RBD or S1 domain variants. Binding of tri-Ty1 was obstructed by the amino acid change E484K, which is found in the Beta VOC and linked to neutralization escape (12, 16, 51) The multimodular nanobodies comprised of modules that target distinct epitopes, tri-TMH and tri-TMV, retained a level of binding to the E484K RBD variant. The fluorescence signals of Dylight 633-labelled trimeric nanobodies tri-Ty1, tri-TMH and tri-TMV bound to the different RBD and S1 mutants in the array were normalized relative to the corresponding wt protein signals in the same array. Error bars represent the standard deviation of two replicate wells. **B**) Location of the amino acid changes within the SARS-CoV-2 RBD in the Alpha and Beta variants. Amino acid change N501Y is found in the Alpha and Beta variants (15, 16), and the Beta variant displays the additional changes K417N and E484K (16).

### Multimodular nanobodies potently neutralize SARS-CoV-2 wild-type and Alpha

We determined the neutralization potency of multimodular nanobodies against SARS-CoV-2 variants *in vitro*, using a plaque-reduction neutralization assay in VeroE6-TMPRSS2 cells (Figure 3). Multimodular nanobodies neutralized wild-type virus (50 pfu) at ultra-high potency, with IC50 values ranging from 160.9 pM for tri-Ty1, to 83.66 pM for tri-TMV and 50.11 pM for tri-TMH. This result shows that multimodular structure improves neutralization efficacy from constituent nanobodies up to thousand-fold. Neutralization was also tested with virus administered at 1 MOI, where the relative efficacies of the nanobodies followed the same trends as in the 50 pfu assay, although at lower IC50-values ranging from 400 pM to 2 nM (SI Figure S5).

**Figure 3.**
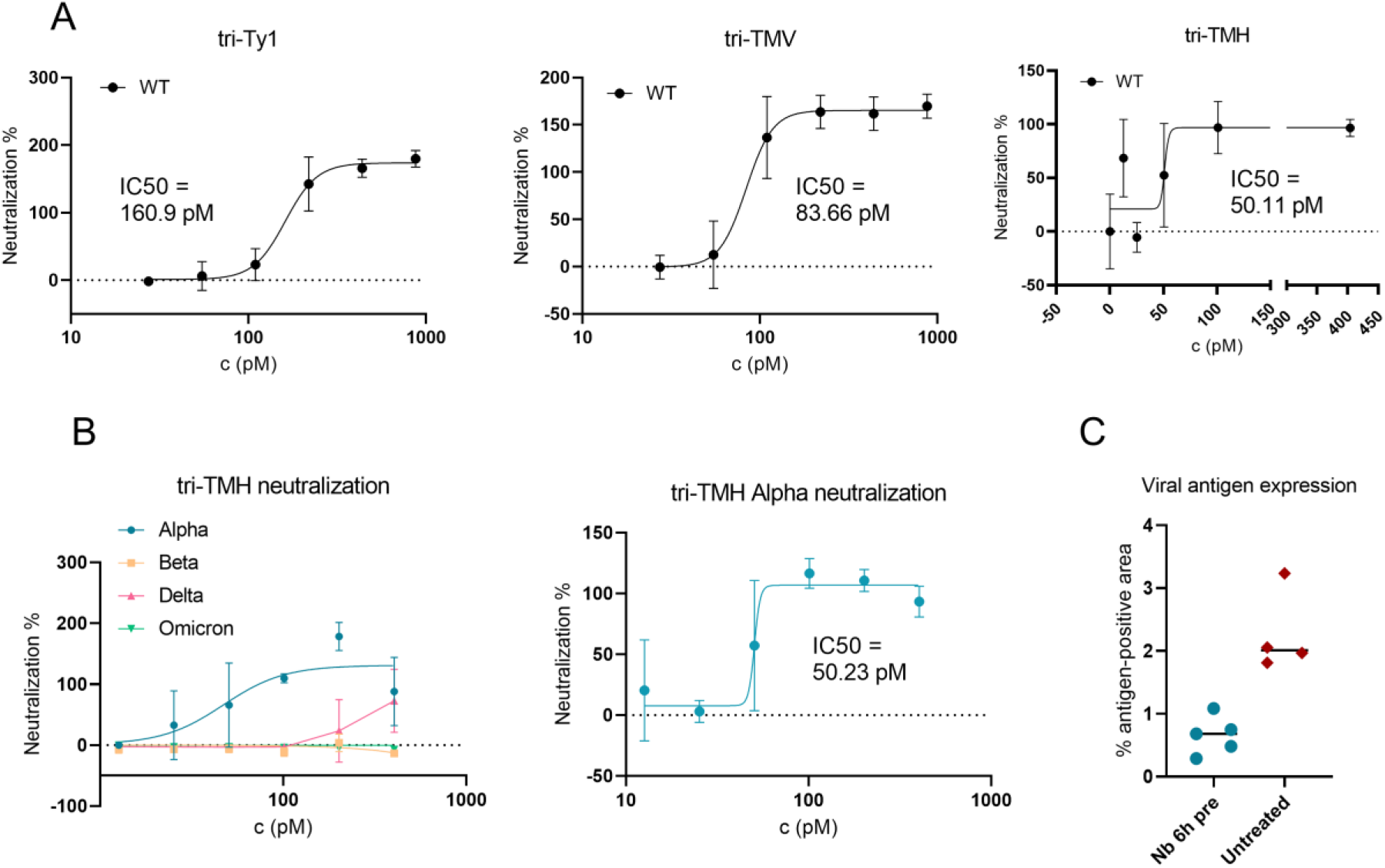
Multimodular nanobodies neutralize SARS-CoV-2 wild-type, Alpha and Delta variants at high potency and limit the course of the disease in a hamster model. **A)** Wild-type SARS-CoV-2 is neutralized by nanobodies tri-Ty1, tri-TMV, and tri-TMH. **B)** Tri-TMH neutralizes the Alpha and Delta variants. **C)** A morphometric analysis of the presence of SARS-CoV-2 antigens in lung tissue sections shows a decrease in viral antigen expression in hamsters pre-treated with multimodular nanobody (Nb 6h pre), compared to untreated individuals (n=5 in nanobody treated group, n=4 in untreated).

To investigate the capability of multimodular nanobodies to neutralize SARS-CoV-2 variants, we tested tri-TMH, the most potent neutralizer of wild-type SARS-CoV-2, against the VOCs Alpha, Beta, Delta, and Omicron. Tri-TMH neutralizes SARS-CoV-2 Alpha variant with equivalent potency to wild-type (IC50 = 50.23 pM), and Delta with decreased potency (estimated IC50 ∼600 pM). Beta and Omicron variants, however, escaped neutralization (Figure 3B). This contrasts with the antigen microarray data (Figure 2), where nanobodies with three distinct modules were found to retain most of their binding to RBD with this mutation, which may be due to the sensitive binding array method detecting weak binding insufficient for neutralization.

The inhibitory effect observed in neutralization assays against SARS-CoV-2 wild-type was recapitulated in an animal model for SARS-CoV-2 infection. We tested the *in vivo* efficacy of the tri-TMH nanobody, the most potent neutralizer identified by our *in vitro* neutralization assays, in 8-week-old Syrian golden hamsters. A 30 ug dose of nanobody was administered to six hamsters intranasally six hours before infection with wild-type (Wuhan-Hu-1) SARS-CoV- 2. After three days, the hamsters were euthanized, and tissue samples taken. SARS-CoV-2 was detected and quantified in the lung tissue of the nanobody group and an untreated control group via RT-qPCR of RdRp and E genes, and via a morphometric immunohistology assay. Both RT-qPCR and morphometric analysis of lung sections revealed a ∼70% reduction of viral antigen positive tissue area and a decrease in viral gene expression in nanobody-treated animals (Figure 3, panel C; Figure S4).

### Cryo-EM analysis reveals the conformational landscape of SARS-CoV-2 spike bound to a multimodular nanobody

While many individual nanobody modules and their binding epitopes on the SARS-CoV-2 spike have been structurally characterized, these reconstructions do not account for the potential steric constraints imposed by linker-bound modules in multimodular nanobodies (29-37, 53). We applied cryo-EM to determine the binding mode of multimodular nanobody tri-TMH to spike.

The natural variation in SARS-CoV-2 spike conformation leads to the presence of distinct subpopulations in cryo-EM data, primarily fully closed (“all down”) and partially open (“1 up”) states (4, 7, 54). While differential distributions of the two states have been reported, native spikes on the viral surface show 31% and 55% of fully closed and partially open conformations, respectively (54). Furthermore, certain neutralizing antibodies have been shown to disrupt the conformation of the spike due to steric incompatibility with the prefusion state (55, 56).

We set out to determine how nanobody tri-TMH impacts the conformation of the spike. Our cryo-EM data show the spike retaining a prefusion conformation, with subpopulations presenting the closed and partially open states. Both states are fully bound with tri-TMH (Figure 4A), although due to the variable placement of the modules, local resolution does not allow the unambiguous identification of individual nanobody moieties. Our results indicate that the inclusion of multiple simultaneously binding modules, linked by (GGGGS)_4_ linkers, does not result in sufficient steric strain to disrupt the prefusion conformation, allowing the native distribution of RBD conformational states. We postulate that the increased potency of multimodular inhibitors is primarily due to enhanced avidity, not altered mechanistic properties.

**Figure 4.**
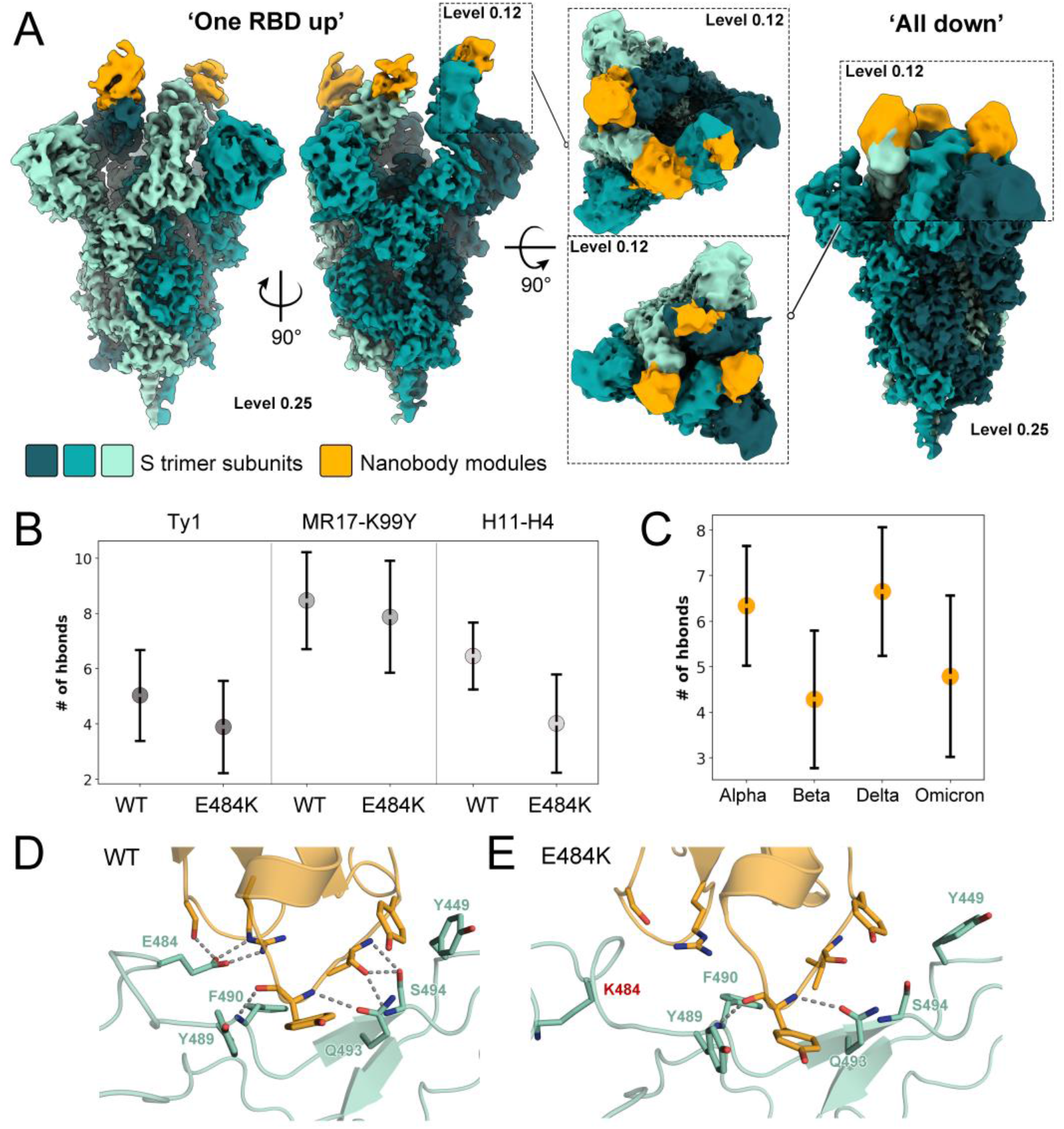
Insights into multimodular nanobody binding from cryo-EM analysis and molecular dynamics simulations. A) Cryo-EM reconstructions of the S-trimer with multimodular nanobody tri-TMH bound. The reconstructions show the typical spike protein conformations (one RBD up and all RBDs down). B) Number of hydrogen bonds between RBD (wild-type or with E484K amino acid change) and each nanobody module of tri-TMH observed from MD simulation data. The dot shows the mean value based on all simulation replicas, and the line represents the standard deviation. C) The number of hydrogen bonds between nanobody module H11-H4 and the RBD of SARS-CoV-2 VOCs, as predicted by MD simulations. D) The interface between nanobody module H11-H4 (orange) and wild-type RBD (green). E) The interface between nanobody module H11-H4 (orange) and RBD (green) with amino acid change E484K (highlighted in red).

### Molecular dynamics simulations indicate re-arrangement of nanobody-RBD interface as a result of amino acid changes present in VOC

Following the cryo-EM analysis, we sought to elucidate the molecular basis for neutralization evasion as observed for VOC Beta (Figure 3) via molecular dynamics (MD) simulations. Interestingly, many nanobodies, as well as antibodies derived from B cells of COVID-19 convalescent and vaccinated individuals, show a salt bridge between an antibody scaffold arginine (R52 in nanobodies) and residue E484 of the RBD (15, 16). As amino acid change E484K is linked to neutralization evasion by VOCs (16, 22, 57), we sought to determine how E484K impacts the interface of RBD and nanobody modules included in tri-TMH. MD simulations of both WT RBD and E484K RBD were performed with the individual nanobody modules.

For modules MR17-K99Y and H11-H4, the R52-E484 salt bridge was identified to be extremely stable throughout the WT simulations and in multiple independent simulation replicas (ca. 95 % of total simulation time). However, when amino acid change E484K was simulated, the interaction to R52 was rapidly broken (within 1 ns). The dissociation of the salt bridge as a consequence of charge change from anionic glutamate to positively charged lysine contributed to higher instability of the E484K RBD, with the surrounding loop becoming more mobile (Figure S2 and S3, Figure 4D-E). In addition, the number of hydrogen bonds between RBD and the nanobody modules decreased in the E484K systems (Figure 4B). Most of these disrupted hydrogen bonds were in the conserved RBD binding epitope (Figure 4D-E). Additional simulations were performed with the full set of VOC mutations in the RBD. Here, the Alpha and Delta VOC, where E484 is preserved, behaved similarly to the WT systems. However, both Beta and Omicron VOC, where E484 is substituted by lysine and alanine respectively, showed higher instability (SI Figure S2) and a decrease in hydrogen bonding interactions throughout the simulation (Figure 4C). These data indicate that E484 is central for strong nanobody binding.

### Nanobodies fused to split nanoluciferase fragments detect SARS-CoV-2 Spike at picomolar concentrations

Here, we applied nanobody engineering in the development of a SARS-CoV-2 detection assay. The modular nature of nanobodies makes them amenable to fusion with signal molecules, and their small size allows the targeting of proximal epitopes, such as those presented by the three subunits of the SARS-CoV-2 spike. These properties align with the principle of protein-fragment complementation assays, where the activity of a split signal molecule is restored once the split fragments are brought into close proximity by the interaction of proteins fused to the fragments (58). Nanobodies shown in Figure 1 target epitopes in the RBD, and we hypothesized that, when fused to split signal molecule fragments, their binding to three spike subunits reconstitutes the signal molecule, allowing sensitive detection in a single step.

We selected the split version of NanoLuc, an engineered 19-kDa luciferase with enhanced stability and brightness (43, 44), as the signal molecule for the assay. RBD-binding nanobody Ty1 (28) was fused with fragments of NanoLuc, titled SmBit or LgBit (42, 44), using flexible (GGGGS)_4_ linkers to connect nanobodies with signal molecule fragments (Figure 5). We performed proof-of-principle experiments to gauge the ability of the nanobody-nanoluciferase fusions to detect recombinantly produced SARS-CoV-2 Spike in solution. 10 nM concentrations of each nanobody fusion, one fused to the LgBit and the other to the SmBit fragment, were mixed with a dilution series of either recombinant spike or negative control protein (bovine serum albumin, BSA) and incubated for 15 minutes before substrate addition and luminescence measurement. The results show that the spike signal can be distinguished from an equimolar concentration of the negative control protein at a concentration as low as 200 pM (Figure 5). These proof-of-principle experiments demonstrate the potential of nanobodies for diagnostic agent development.

**Figure 5.**
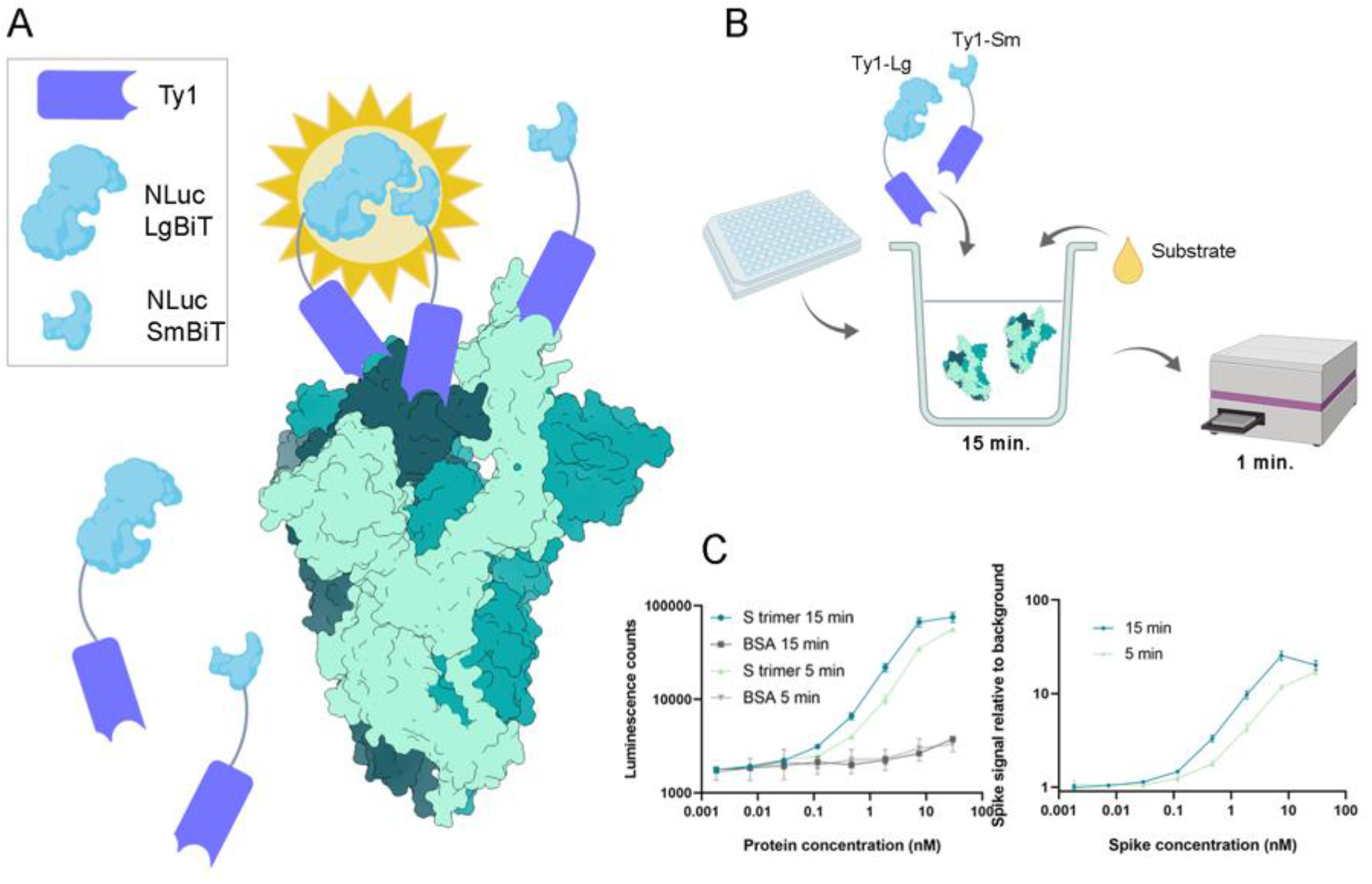
SARS-CoV-2 spike protein detection with a proximity-triggered assay. A) Split nanoluciferase fragments are fused to Ty1 nanobody (28) with GGGGS4 linkers. The enzymatic activity of the luciferase is restored upon nanobody binding to adjacent RBDs. B) The reaction setup. The nanobody fusions are added into the wells of an opaque white plate with recombinant spike, and the mixture is incubated for 15 minutes. Then, the substrate is added, and the luminescence reads are recorded from each well. C) and D) The signal caused by spike protein can be distinguished from background down to spike concentration of 200 pM.

## Discussion

In this study, we created potent SARS-CoV-2 neutralizers and a detection assay concept through fusions of spike-targeted nanobody modules. Our multimodular nanobodies were linked together with (GGGGS)_4_-linkers to facilitate simultaneous binding to multiple epitopes and to increase the avidity of binding. We observed a notable increase (up to 1000-fold) in the neutralization potency of the multimodular nanobodies relative to the reported IC50 values of the single constituent nanobodies (SI Table S1) in neutralization assays, with SARS-CoV-2 Alpha and Delta VOCs being neutralized, but not Beta and Omicron. This observation is in line with recent works reporting nanobody multimerization increasing neutralization potency (38, 41, 59, 60). Multimodular nanobodies showed IC50 values in the 50−160 pM range (Figure 3A-B) in cell culture, and, in an animal model, prophylactic administration of tri-TMH in the nasal cavity limited tissue damage in the lungs (Figure 3C).

Our MD simulation results indicate that the E484K amino acid change results in a loss of salt bridge with conserved R52 and conformational rearrangement in the receptor-binding domain (Figure 4), causing disruption of the nanobody binding interface and leading to neutralization escape in Beta and Omicron variants. MD simulation data also revealed interaction trends (Fig 4, panels D and E) at the antigen interface that correlate with the experimentally observed neutralization (Figure 3). We postulate that this approach, focused on simulating hydrogen bonding networks and RMSD trends at the antigen interface may be applied to qualitatively predict the potential for escape from specific neutralizers.

Drawing from protein-fragment complementation assays, we were inspired to combine nanobody modules with signal molecules to develop new diagnostic tools. Our nanobody-based assay was successful at detecting low concentrations of SARS-CoV-2 spike protein. This is in line with the limit of detection reported for other antigen tests, such as a FRET-based assay (61) and similar to commercially available rapid point-of-care antigen tests (40). To our knowledge, this is the first report of a nanobody-based detection approach that makes use of the protein-fragment complementation toolkit. In contrast to monoclonal antibodies commonly used in diagnostics, engineered nanobodies have multiple attractive properties, including cheap and scalable production that does not require resource-intensive mammalian cell culture (48). It is also becoming feasible to discover nanobodies from synthetic libraries, allowing the transition away from conventional animal immunization prevalent in antibody generation (62). We envision that this combination of nanobody modules with split signal molecules presents a powerful platform for the rapid development of single-step detection assays for emerging pathogens.

Emerging SARS-CoV-2 variants continue to challenge the efficacy of therapeutic strategies. Nanobodies, being inexpensive and readily modified, show promise as inhibitor candidates and diagnostic tools. Our work showcases the potential of nanobodies engineered to target viral pathogens to improve preparedness for outbreaks of emerging infectious disease.

## Materials and Methods

### Production and purification of multimodular and luciferase-fused nanobodies

Synthetic genes encoding multimodular nanobodies (individual modules reported in (28-30, 41)) and nanobody fusions with split nanoluciferase (44, 45) were ordered cloned in expression vector pET100/D-TOPO (GeneArt, Thermo Fisher Scientific). Nanobodies were expressed in *Escherichia coli* Rosetta-gami 2 (DE3) cells (Novagen) in autoinduction media and purified through consecutive nickel affinity chromatography and size-exclusion chromatography (see Supplementary Methods).

### Production and purification of recombinant SARS-CoV-2 S protein

SARS-CoV-2 spike was expressed from a synthetic cDNA template (GeneArt, Life Technologies) encoding the S protein ectodomain residues 14−1208 from the Wuhan-Hu-1 strain (NCBI Reference Sequence: YP_009724390.1) stabilized in the prefusion state (4, 7) with proline substitutions at residues 986 and 987, an abrogated furin S1/S2 cleavage site with a “GSAS” substitution at residues 682–685, and a C-terminal T4 fibritin trimerization motif. In our construct, the trimerization motif was followed by an HRV3C protease cleavage site, SpyTag003 (63), and 8xHisTag. The gene was cloned into the mammalian expression vector pHLsec (64) and transfected into Expi293F™ (Thermo Fisher Scientific) suspension cells for expression. The protein was purified through nickel affinity chromatography (see Supplementary Methods for further details).

### Cryo-EM grid preparation, data acquisition and data processing

A 3-μl aliquot of a pure, prefusion SARS-CoV-2 S-trimer (0.3 mg/ml) mixed with tri-TMH (0.05 mg/ml) was applied on Quantifoil 1.2/1.3 grids (1.2-μm hole diameter, 200 mesh copper) that had been glow discharged in Plasma Cleaner PDC-002-CE (Harrick Plasma) for 30 s. The grids were blotted for 6 s and plunged into liquid ethane using a vitrification apparatus (Vitrobot, Thermo Fisher Scientific).

Data were collected on a Titan Krios transmission electron microscope (Thermo Fisher Scientific) equipped with Gatan K2 direct electron detector. EPU v 2.11.0 software was used to acquire micrographs, and images were collected with a dose of 1.38 e⁻/Å² per image (SI Table S2).

Data were processed in cryoSPARC (65). Movie frames were aligned and averaged to correct for beam induced motion. Contrast transfer function (CTF) parameters were estimated using CTFFIND4 (66). An initial set of particles, picked with the blob-picker, was classified and the particles in good 2D classes were used to train Topaz particle picker (67, 68). A total of 91,601 particles were selected after cleaning the picked set with 2D classification. An initial volume with C3 symmetry was calculated *ab initio*. The volumes were refined following a strategy described in Supplementary Methods.

### Model fitting into cryo-EM maps

Molecular models of the S trimer (PDB: 7A29) and nanobody-RBD complexes (PDB: 6ZHD, 6ZXN, 7CAN) were fitted in the cryo-EM density map with the “fitmap” function in USCF Chimera (69) (for a detailed description, see Supplementary Methods).

### Proof-of-concept tests for the detection assay with Split NanoLuc-nanobody fusions

A triplicate dilution series of purified recombinant Spike in Tris buffer (10 mM Tris-HCl pH 7.5, 150 mM NaCl) was made on an opaque white 96-well plate. In each well, purified Ty1-LgBiT and Ty1-SmBiT were added at 10 nM final concentration. The plate was incubated for 15 minutes at room temperature, after which nanoluciferase substrate coelenterazine H was added to 200 nM concentration. Luminescence readings were measured with Perkin-Elmer EnSpire multimode plate reader using the ‘Luminescence’ program. To determine the signal-to-noise ratio in the luminescence reaction, the assay was performed on an equimolar dilution series of BSA. To calculate the final luminescence measurement while taking noise into account, the average readings of each triplicate sample were divided by corresponding averages from the BSA controls.

### Antigen array

Nanobodies were labelled with DyLight 633 in PBS supplemented with 50 mM sodium borate (pH 8.5) at 1 mg/mL protein concentration and 50 µM Dylight 633 NHS ester (Thermo Fischer Scientific) at room temperature for 2h, followed by removal of unreacted dye with Zeba Spin 7K MWCO desalting columns (Thermo Scientific).

Wild type and variant SARS-CoV-2 RBD and Spike S1 domains were biotinylated and arrayed as duplicate spots (0.1 ng per spot) in the wells of streptavidin-coated microtitration plates using a piezoelectric non-contact microarray dispenser (Nano-Plotter, GeSiM, Germany). The antigens were purchased from the following sources: RBD wt (aa 319-541 of the S protein) and S1 wt (aa 14-681 of the S protein) from Medix Biochemica; RBD single mutants K417N, E484K and N501Y and S1(K417N, E484K, N501Y, D614G) quadruple mutant from SinoBiological.

The antigen arrays in microplate wells were blocked with 50 µL of Assay buffer (Tris-buffered saline (TBS), pH 8.0 + 0.05% Tween 20) per well for 30 min at RT, followed by three washes with washing buffer (TBS containing 0.05% Tween 20). DyLight 633 -labelled nanobody solutions (1 µg/mL in Assay buffer) were added 50 µL/well, incubated in a plate shaker at 600 rpm, RT for 1h, followed by three washes with washing buffer. Residual liquid droplets were removed by centrifuging the plate upside down on a paper towel in a plate adapter (453 g, 1 min), after which the plate was let dry for 15 min in a 37°C room. The Dylight 633 -labelled nanobodies bound to the arrayed antigens were detected by fluorescence scanning through the clear bottom of the microplate with a Tecan LS400 confocal laser scanner, using a 633 nm laser for excitation and a 670/25 emission filter.

The fluorescence scan images were analyzed with Array-Pro Analyzer software (Media Cybernetics) and the raw spot signal data was exported to Microsoft Excel for further calculations. Net signals were obtained by subtracting the well background from the raw spot signals (average pixel intensity in the spot area), after which the spot signals were normalized to the wild-type antigen spot signals in the same well: single RBD mutant signals were expressed as percentage of the RBD wt signal whereas the quadruple S1 mutant signal was expressed as percentage of the S1 wt signal.

### Neutralization assays

VeroE6-TMPRSS2-H10 cells (61) were seeded to 96-well plates (white-sided optically clear bottom PerkinElmer) in density 30 000 cells/well 24 h before the assay. The nanobodies were diluted in series 1:150 000, 1:300 000, 1:600 000, 1:1 200 000, 1:1 240 000, and 1:4 800 000 in virus growth medium (VGM) containing MEM (Sigma, 2279), 2 % FBS, L-glutamine, and 1x penicillin-streptomycin. Diluted nanobodies were mixed with 50 pfu (Figure 3) or MOI 1 (SI Figure S5) (for wild-type, Alpha, Beta, and Delta) or MOI 0.2 (for Omicron) virus and incubated for 1 h at 37°C 5% CO_2_. Thereafter, cells were treated with the mixture of nanobodies and virus. The virus dilution in VGM without nanobodies was used as a negative control and non-infected cells (MOCK) was used as a positive control of the nanobody neutralization effectiveness. After 5 days of incubation the medium was removed, and cells were treated with CellTiter-Glo 2.0 cell viability assay reagent (Promega, G9243) for 20 min at RT. Then, the cellular ATP was measured via the detection of luminescent signal using HIDEX Sence microplate reader (Hidex Oy, Finland) Viability of the MOCK infected cells was considered as 100 %. Neutralization efficacy percentage for each measurement was calculated considering MOCK infected cells as 100% neutralization and untreated, virus infected cells as 0% neutralization. Curve-fitting and IC50 calculation was performed for the normalized neutralization data with GraphPad Prism version 9.2.0 for Windows, using the Nonlinear regression method (Absolute IC50, X is concentration), with baseline constraint set to zero.

### Animal experiments

A total of 20 Syrian Golden hamsters (Scanbur, Karl Sloanestran, Denmark) were moved to the University of Helsinki biosafety level-3 facility and allowed to acclimatize to individually ventilated biocontainment cages (ISOcage; Scanbur, Karl Sloanestran, Denmark) for seven days with ad libitum water and food (rodent pellets) prior to infection.

For the main experiment, six 8-week-old male and female Syrian Golden hamsters received 30 µg of nanobody, in 0.61 mg/mL concentration, 6 h prior to intranasal infection with 5 × 10^4^ SARS-CoV-2 (wt/D614G strain). The control group received an equal volume of PBS (n=4).

Euthanasia was performed under terminal isoflurane anaesthesia with cervical dislocation, followed by dissection immediately after death. Samples were collected from the lungs for RT-qPCR (see Supplementary Methods), and the remaining lung tissue, including trachea, heart, oesophagus, and bronchial lymph nodes, was immersed in 10% buffered formalin. After 48h, the tissue was stored in 70% ethanol until processing for histological and immunohistological examination.

### Histological, immunohistological and morphometrical analyses

Three to five cross sections were prepared from the fixed lung tissue and routinely paraffin wax embedded. Consecutive sections (3-5 µm) were prepared and stained with hematoxylin eosin (HE) for histological examination or subjected to immunohistological staining for SARS-CoV-2 antigen expression, using a previously published staining protocol (70). For immunohistology, the horseradish peroxidase method was applied. Rabbit anti-SARS-CoV nucleocapsid protein (Rockland, 200-402-A50) served as the primary antibody, and DAB (EnVision FLEX DAB+ Chromogen in Substrate buffer; Agilent) for visualization of antibody binding. All incubations took place in an autostainer (Dako). Sections were subsequently counterstained with haematoxylin.

For morphometric analysis, the immunostained sections were scanned (NanoZoomer-XR C12000; Hamamatsu, Hamamatsu City, Japan) and analysed using the software programme Visiopharm (Visiopharm 2020.08.1.8403; Visiopharm, Hoersholm, Denmark) to quantify the area of viral antigen expression in relation to the total area (area occupied by lung parenchyma) in the sections. This was used to compare the extent of viral antigen expression in the lungs between untreated and treated animals. A first app was applied that outlined the entire lung tissue as ROI (total area). For this a Decision Forest method was used and the software was trained to detect the lung tissue section (total area). Once the lung section was outlined as ROI the lumen of large bronchi and vessels was manually excluded from the ROI. Subsequently, a second app with Decision Forest method was trained to detect viral antigen expression (as brown DAB precipitate) within the ROI.

### Molecular dynamics simulations of nanobody-RBD interface

MD simulations were performed on the model systems constructed using the cryo-EM structure of nanobody monomers and RBD (PDB 6ZXN, 7CAN, 6ZBP). During model system construction long glycine linkers were removed. VMD psfgen tool (71) was employed to add missing hydrogen atoms, make point mutations, and solvate the systems with TIP3P water and 0.1 M NaCl to give a total system size of ca. 150,000 atoms. The force field for all components of the system was CHARMM36 (72). GROMACS v20.3 (73) was used for all equilibration and production simulations. First, the initial systems were minimized with restraints on heavy protein atoms (20,000 kJ mol^−1^ nm^−2^). Following convergence, a 100 ps NVT equilibration was performed with the same restraints, followed by a 10 ns NPT with the restraints only on the protein backbone atoms. All restraints were removed for the production runs, which employed the LINCS algorithm (74) to achieve a 2 fs timestep and used the Nosé -Hoover thermostat (75, 76) to maintain 310 K temperature and the Parrinello-Rahman barostat (77) to maintain 1 atm pressure. The electrostatic interactions were controlled by the particle mesh Ewald method (78) with a 12 Å cutoff, while the van der Waals cutoff was also 12 Å, with a switching distance of 10 Å. Multiple independent simulation replicas were performed and trajectories were visualized and analyzed using VMD (71) and Pymol (The PyMOL Molecular Graphics System, Version 1.2r3pre, Schrödinger, LLC). Table S3 lists system setups and their respective lengths.

### Molecular graphics

Molecular graphics images were generated using PyMOL (The PyMOL Molecular Graphics System, Version 2.4.0a0, Schrödinger, LLC), UCSF Chimera (69), and UCSF ChimeraX (79).

### Data accession

Cryo-EM reconstructions of the nanobody-bound SARS-CoV-2 spike in the one-up and all-down states have been deposited at EMDB under the accession codes EMD-XXXX and EMD- YYYY.

## Acknowledgements

This work was funded by the Academy of Finland (project number 336492/JH, 336490/OV, 339510/TS, 338176/VS, 336495/PS), VEO - European Union’s Horizon 2020 (grant number 874735 to OV), the Jane and Aatos Erkko Foundation (to OV, VS), Sigrid Jusélius Foundation (to OV, VS), and Helsinki University Hospital Funds (TYH2018322 and TYH2021343 to OV). The authors acknowledge CSC – IT Center for Science, Finland, for computational resources. Cryo-EM data was collected at the Umeå Core Facility for Electron Microscopy, a node of the Cryo-EM Swedish National Facility, funded by the Knut and Alice Wallenberg, Family Erling Persson and Kempe Foundations, SciLifeLab, Stockholm University and Umeå University. The use of the facilities and expertise of the Instruct-HiLIFE cryo-EM unit, member of Biocenter Finland and Instruct-FI, is gratefully acknowledged, as well as the staff at HUSLAB Virology and Immunology for providing samples for virus isolation. Part of the work was carried out with the support of HiLIFE Laboratory Animal Centre Core Facility, University of Helsinki, Finland.

