## Supplementary information for "Nanobody engineering for SARS-CoV-2 neutralization and detection"

**Items:**

**Figure S1**. Distances between key residues in the receptor-binding domains in the different conformations of the SARS-CoV-2 spike

**Figure S2.** RMSD (Root Mean Square Displacement) of protein Cα atoms over MD simulation

**Figure S3**. Distance between charged residues (E/K and R) in MD simulations

**Figure S4**. Nanobody treatment leads to decreased viral replication in hamster lungs

**Figure S5.** Neutralization of SARS-CoV-2 WT at 1 MOI by multimodular nanobodies

**Figure S6.** Resolution estimates of cryo-EM reconstructions

**Figure S7.** Whole cryo-EM maps

**Figure S8.** Fitting of PDB models in the cryo-EM maps

**Table S1.** Reported binding and neutralization properties of nanobody modules

**Table S2.** Cryo-EM data collection and processing statistics

**Table S3.** Parameters for the molecular dynamics simulations

**Supplementary Methods.**

**Supplementary References.**

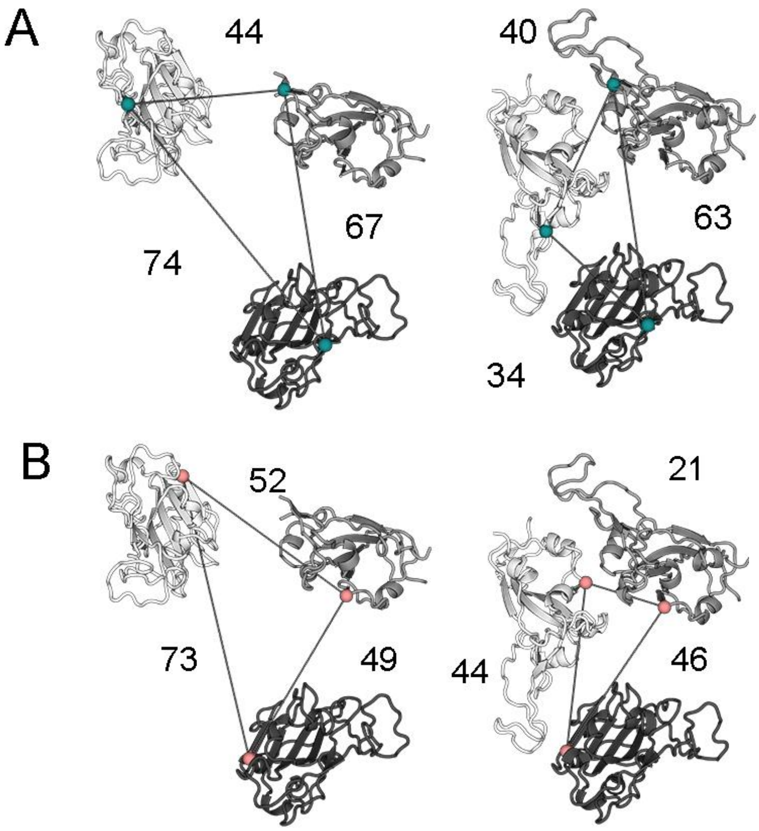


**Figure S1.** Distances (Å) between key residues in the receptor-binding domains in the different conformations of the SARS-CoV-2 spike, illustrating the distance bridged by nanobody modules connected by 20-AA linkers. A) Distances between the central residues (493) of the ACE2 binding sites in the 3 RBDs, with spike in the 2-up conformation (left) or 1-up conformation (right). B) Distances between the central residues (375) of the VHH V nanobody epitopes in the 3 RBDs, with spike in the 2-up conformation (left) or the 1-up conformation (right).

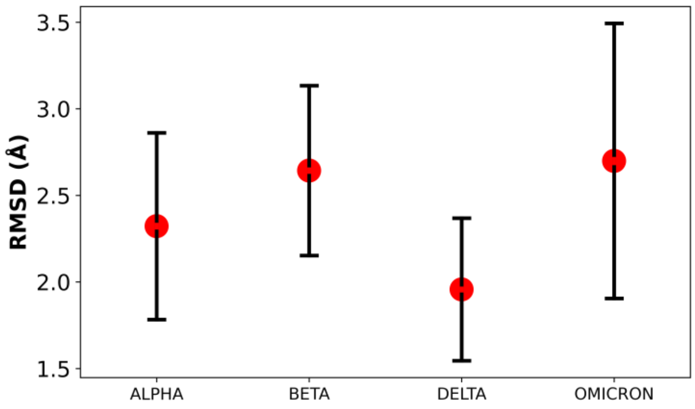


**Figure S2.** RMSD (Root Mean Square Displacement) of protein Cα atoms over MD simulation using the zero frame as a reference. Data from all simulation replicas are combined, and the mean value is shown as a dot, with the standard deviation as error bars.

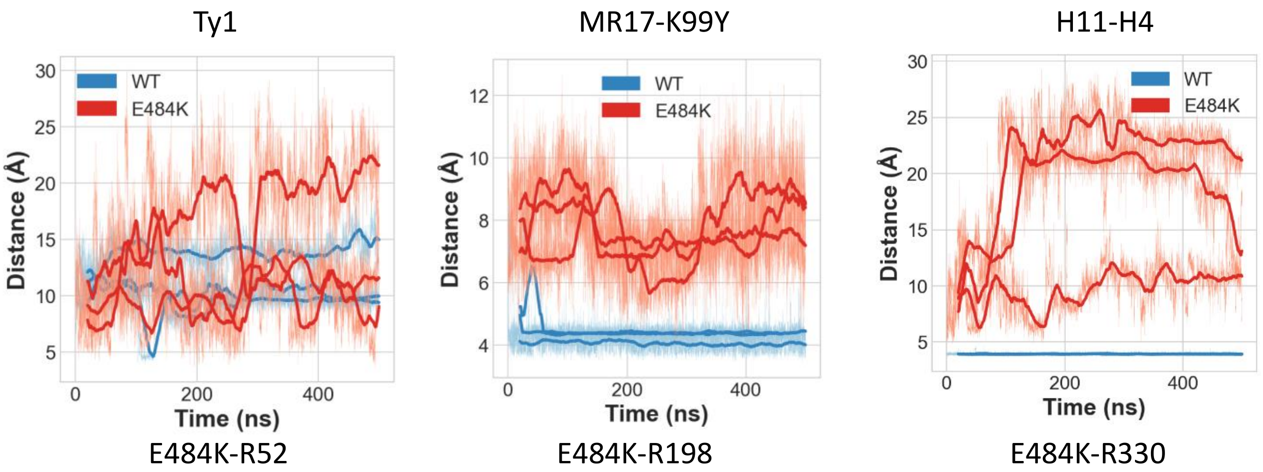


**Figure S3.** Distance between charged residues (E/K and R) in MD simulations. The three simulation replicas are shown separately for WT (blue) and E484K (red). The bold lines represent a running average of the previous 20 ns simulation data. Distances were measured between the glutamic acid CD atoms or lysine NZ atoms, and arginine CZ atoms.

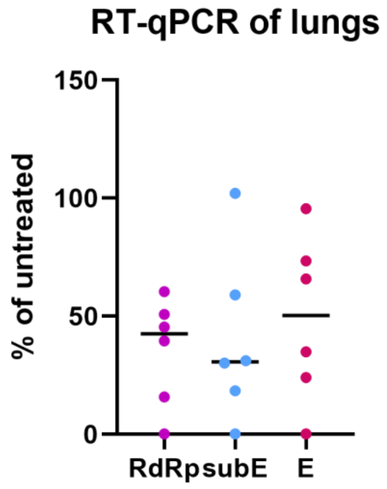


**Figure S4.** RT-qPCR of hamster lung samples. Results shown as percentages of the average value for untreated individuals (n=4).

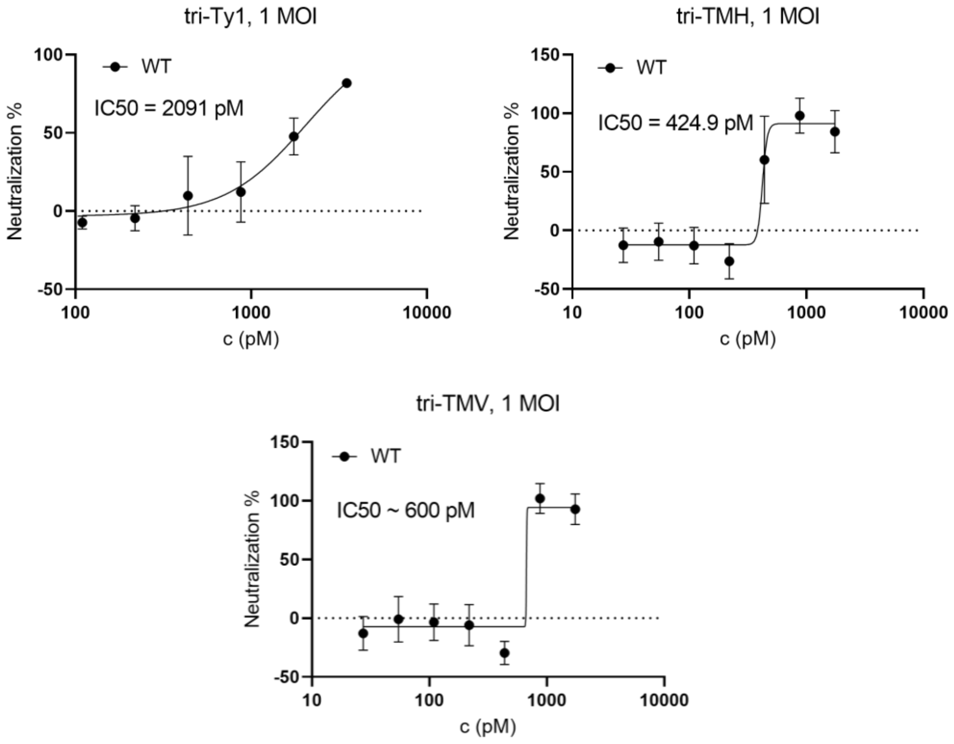


**Figure S5**. Parallel neutralization experiments were performed with 1 MOI of virus instead of 50 pfu (Main Figure 3). Nanobodies tri-Ty1, tri-TMH, and tri-TMV show neutralization at these higher virus titers, but the calculated IC50 values are decreased. With higher amounts of virus used for infection, tri-TMH remains the most effective neutralizer, based on the calculated IC50 value.

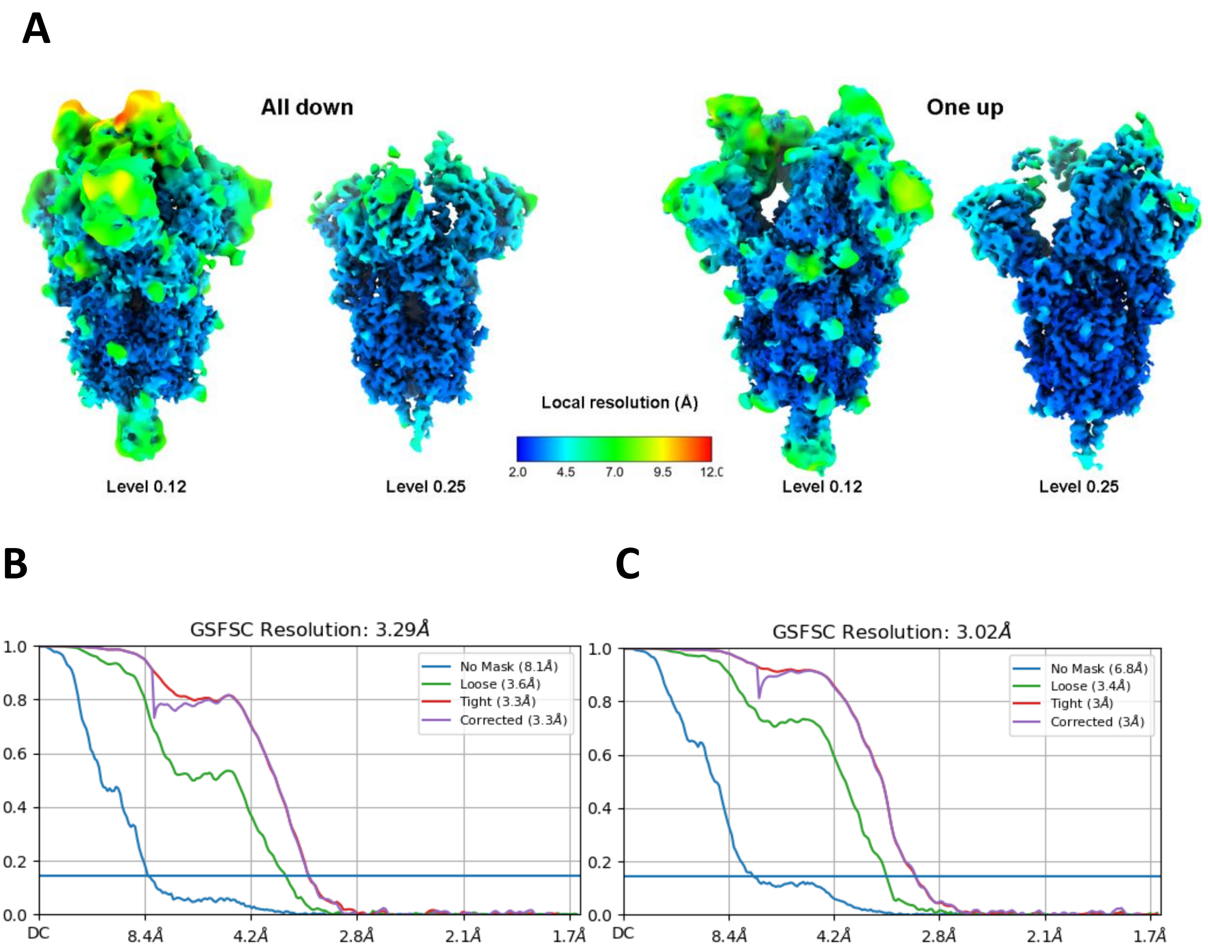


**Figure S6. Resolution estimates of cryo-EM reconstructions. A)** Local resolution was estimated and the maps were filtered in cryoSPARC. The one-up and all-down maps are each shown at two volume levels. **B-C)** FSC curves for the all-down map (B) and the one-up map (C).

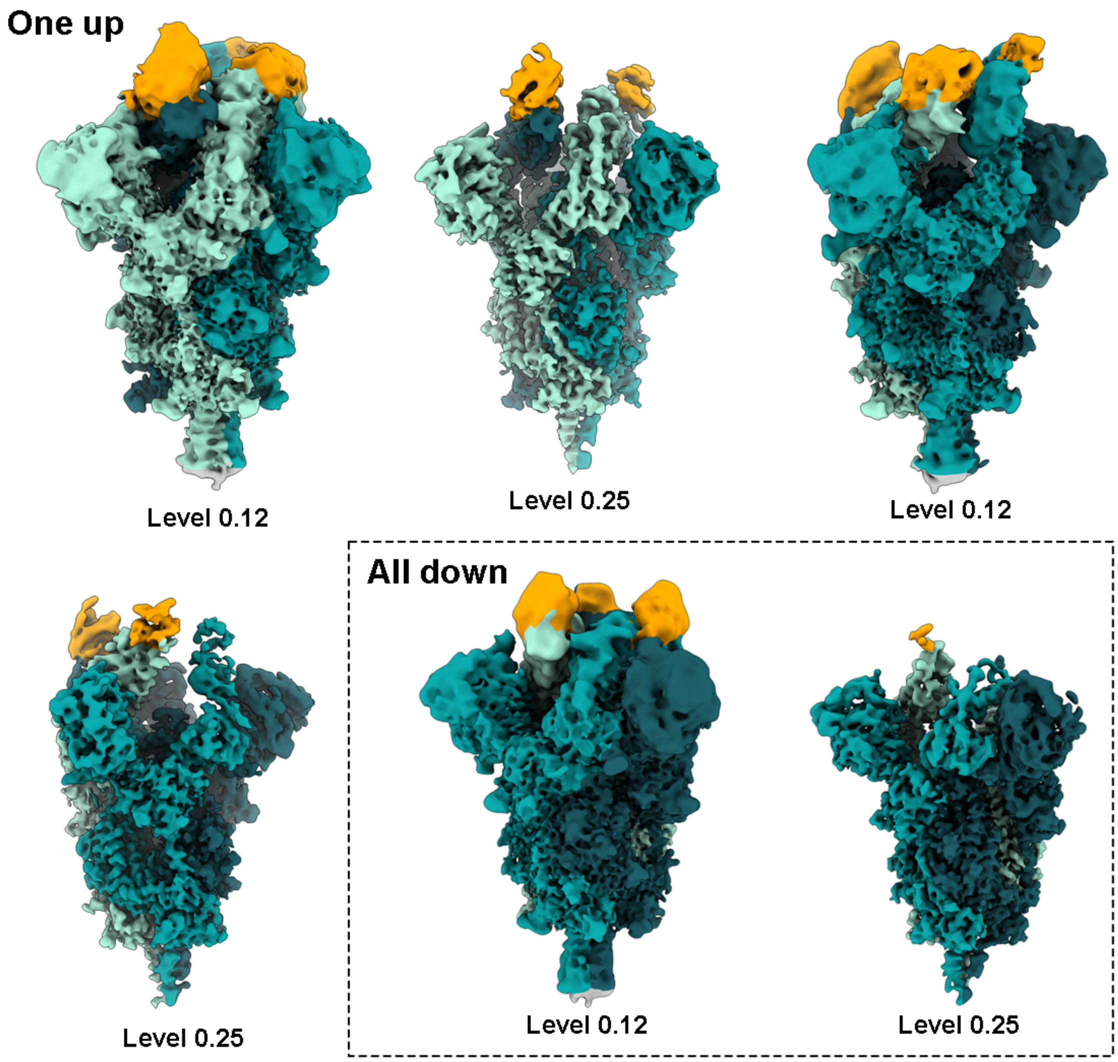


**Figure S7. Whole cryo-EM maps.** The one-up map (two views of the same map) and the all-down map are each shown at two different volume levels. Level 0.12 allows detection of all three nanobodies, whereas level 0.25 filters out flexible regions of the molecule but shows density features in high resolution regions.

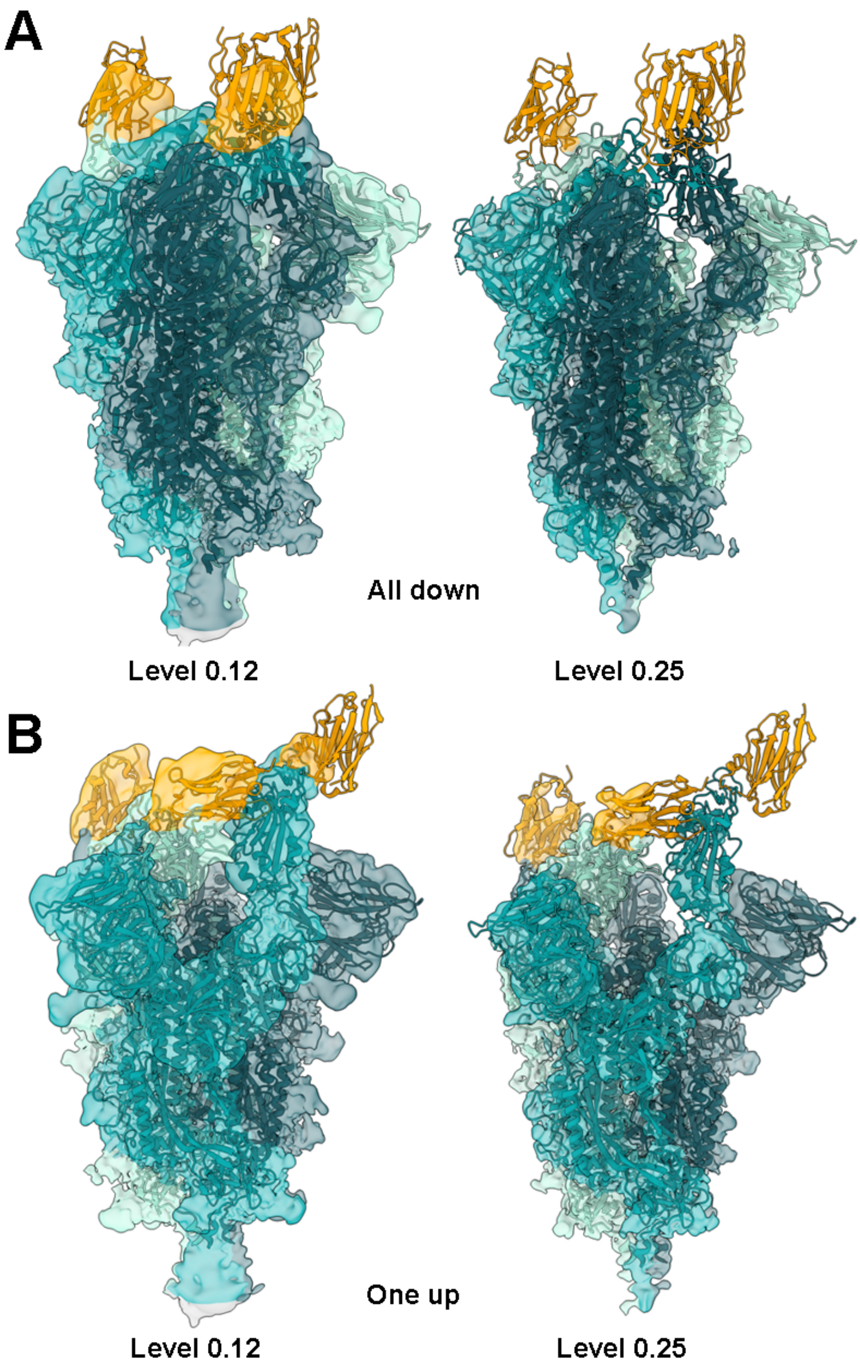


**Figure S8. Fitting of PDB models in the cryo-EM maps.** The S trimer model (PDB: 7A29) and models of nanobody-RBD complexes (PDB: 6ZHD, 6ZXN, 7CAN), displayed as ribbon representations, were fitted into the cryo-EM maps as shown. **A)** Map with all RBDs down. **B)** Map with one RBD up.

**Table S1.** Reported binding and neutralization properties of nanobody modules.

| Nanobody | Citation | $K_D$ (nM) | IC50 (nM) |
|---|---|---|---|
| Ty1 | Hanke et al., 2020 | 5—10 | 54 |
| H11-H4 | Huo et al., 2020 | 12±1,5 | 6* |
| MR17-K99Y | Li et al., 2021 | 33 | 38** |
| VHH V | Koenig et al., 2021 | 8,92 | 142 |

*Neutralization efficiency reported for bivalent construct with human IgG Fc.

**Calculated based on the IC50 value of 0.50 µg/mL and the molecular weight of 13.9 kDa.

**Table S2.** Cryo-EM data collection and processing statistics

| | all-down map (J285) | 1-up map (J275) |
|---|---|---|
| Magnification | 165 000x | 165 000x |
| Voltage (kV) | 300 | 300 |
| Electron exposure (e-/Å$^2$) | 55.0 | 55.0 |
| Defocus range (µm) | -1.5 to -3.0 | -1.5 to -3.0 |
| Pixel size (Å) | 0.82 | 0.82 |
| Symmetry imposed | None | None |
| Initial particle images (no.) | 84049 | 84049 |
| Final particle images (no.) | 26708 | 42440 |
| Map resolution (Å) | 3.29 | 3.02 |
| FSC threshold | 0.143 | 0.143 |

**Table S3.** Description of model systems and MD simulation time scales. Three replicas of 500 nanosecond simulations were run for each simulation setup.

| **Simulation setup n:o** | **Nanobody monomer** | **Mutations** |
|---|---|---|
| **1** | Ty1 | - |
| **2** | Ty1 | E484K |
| **3** | MR17-K99Y | - |
| **4** | MR17-K99Y | E484K |
| **5** | H11-H4 | - |
| **6** | H11-H4 | E484K |
| **8** | H11-H4 | **Alpha**: N501Y |
| **10** | H11-H4 | **Beta**: K417N, E484K, N501Y |
| **12** | H11-H4 | **Delta**: T478K, L452R |
| **14** | H11-H4 | **Omicron**: G339D, S371L, S373P, S375F, K417N, N440K, G446S, S477N, T478K, E484A, Q494R, G496S, Q498R, N501Y, Y505H |

### Supplementary methods

**Expression and purification of multimodular and luciferase-fused nanobodies**. Nanobody expression cultures of *Escherichia coli* Rosetta-gami 2 (DE3) cells (Novagen) were grown at 37 °C) in autoinduction media until OD600 reached 0.5, after which the incubation temperature was lowered to 28 °C. The cells were harvested after 24 h from inoculation. To purify the proteins, bacterial cell pellets were resuspended in lysis buffer (10 mM Tris-HCl pH 7.5, 150 mM NaCl) with protease inhibitors and 10 mM imidazole. Cells were lysed with Emulsiflex C3, and the lysate was clarified by centrifugation at 38 000 × g for 30 min at 4 °C. Proteins were purified by immobilized nickel affinity chromatography with a 5 ml HisTrap FF crude column (Cytiva), using 300 mM imidazole for elution. Concentrated eluates were further purified by size-exclusion chromatography (SEC), using the ÄKTA Go system and a Superdex 75 Increase 10/300 column (Cytiva) in PBS buffer. For neutralization assays, 6xHis-tags were removed by enterokinase cleavage (Bovine enterokinase, GenScript). The cleaved tags and enzyme were removed from the sample using HisPur Ni-NTA resin (Thermo Scientific) or SEC as described above.

**Expression and purification of recombinant SARS-CoV-2 S protein.** The Expi293F™ (Thermo Fisher Scientific) suspension cells were grown at a density of $3 \times 10^6$ cells per ml using the ExpiFectamine™ 293 Transfection Kit (Thermo Fisher Scientific). Transfected cells were cultivated on an orbital shaker at 36.5 °C and 5% CO2 for six days, after which supernatant was harvested, clarified by centrifugation, filtered through a 0.45 µM filter, and supplemented with imidazole to 3 mM final concentration. SARS-CoV-2 S-protein was purified from the supernatant by immobilized nickel affinity chromatography with a 1-ml HisTrap excel column (Cytiva) using 300 mM imidazole for elution. The eluate was concentrated and buffer exchanged to 10 mM Tris pH 8 + 150 mM NaCl buffer, and S-trimer was used for cryo-EM grid preparation immediately after purification.

**RT-qPCR.** RNA was extracted from lung samples using Trizol (Thermo Scientific) according to the manufacturers' instructions. Isolated RNA was directly subjected to one-step RT-qPCR analysis based on a previously described protocol for RdRp (1) and for E and subE genes (2) with TaqMan fast virus 1-step master mix (Thermo Scientific) using AriaMx instrumentation (Agilent, Santa Clara, CA, USA). The actin RT-qPCR used for normalization is described in (3). Fold differences between samples were calculated by the comparative Ct method (4) using the average of normalized Ct values from non-nanobody treated infected animal lung tissues as reference.

**Refinement of cryo-EM maps.** Following generation of an *ab initio* volume with C3 symmetry, a consensus map of the S trimer, with C3 symmetry applied, was resolved to 2.66-Å resolution. Visual inspection showed that the RBD region was poorly defined in the consensus map. 3D variance analysis of symmetry-expanded particles was run using a spherical mask defining the RBD region, six principal modes (i.e. eigenvectors of the 3D covariance) and eight classes (or clusters). Particles in each class were subjected to local asymmetric refinement (standard deviation over the prior of rotations and shifts were 5 degrees and 5 Å, respectively, centered at the box center). This local refinement prevented symmetry expanded particles from rotating over their symmetry copy. Particles in the cluster corresponding to the "all-down" conformation were subjected to a second round of 3D variance analysis using 4 principal modes and 4 classes. Particles from 3 of the 4 classes were combined, symmetry copies removed with the "remove duplicates" function and the "all-down" map was locally refined with C3 symmetry. Resolution of the maps was estimated based on the gold-standard Fourier shell correlation (FSC) criterion of 0.143 (5), and the final maps were filtered to local resolution.

**Model fitting into cryo-EM maps.** To fit molecular models of the S trimer (PDB: 7A29) and nanobody-RBD complexes (PDB: 6ZHD, 6ZXN, 7CAN) in the cryo-EM density map , “fitmap” and “matchmaker” functions were used in USCF Chimera (6). The “fitmap” function was used to simulate a density for each PDB model to the global resolution of each map (3.29Å for the all-down map, 3.02Å for the one-up map). The S trimer model was placed first, yielding placements with map-to-map correlation scores of 0.7269 for the 1-up map and 0.7367 for the all-down map. The “matchmaker” function in UCSF Chimera was used to match models of RBDs with the RBDs of the full spike model. To account for the mobility of the RBDs, the individual RBD models were further fitted into the density by the “fitmap” function (correlation scores for RBD models: 1-up map 0.8064, 0.7843, 0.7824; all-down map 0.7937, 0.7799, 0.8017). Finally, RBD-Nb models (PDB: 6ZHD, 6ZXN, 7CAN) were superposed to the fitted RBDs with the matchmaker function.
